# Dodecaploid *Xenopus longipes* provides insight into the emergence of size scaling relationships during development

**DOI:** 10.1101/2022.08.16.504201

**Authors:** Kelly Miller, Clotilde Cadart, Rebecca Heald

## Abstract

Genome and cell size are strongly correlated across species^1–6^ and influence physiological traits like developmental rate^7–12^. Although size scaling features such as the nuclear-cytoplasmic (N/C) ratio are precisely maintained in adult tissues^13^, it is unclear when during embryonic development size scaling relationships are established. Frogs of the genus *Xenopus* provide a model to investigate this question, since 29 extant *Xenopus* species vary in ploidy from 2 to 12 copies (n) of the genome, ranging from 20 to 108 chromosomes^14,15^. The most widely studied species, *X. laevis* (4n=36) and *X. tropicalis* (2n=20), scale at all levels from body size to cellular and subcellular^16^. Paradoxically, the rare, critically endangered dodecaploid (2n=108) *X. longipes* is a small frog^15,17^. We observed that despite some morphological differences, *X. longipes* and *X. laevis* embryogenesis occurred with similar timing, with genome to cell size scaling emerging at the swimming tadpole stage. Across the three species, cell size was determined primarily by egg size, while nuclear size correlated with genome size during embryogenesis, resulting in different nuclear-cytoplasmic ratios at the mid-blastula transition. At the subcellular level, nuclear size correlated more strongly with genome size, whereas mitotic spindle size scaled with cell size. Our cross-species study indicates that scaling of cell size to ploidy is not due to abrupt changes in cell division timing, that different size scaling regimes occur during embryogenesis, and that the developmental program of *Xenopus* is remarkably consistent across a wide range of genome and egg sizes.

**ONE SENTENCE SUMMARY:** Comparison of size metrics in embryos from different ploidy *Xenopus* species, including the dodecaploid *X. longipes*, reveals distinct size scaling regimes during development.

## RESULTS AND DISCUSSION

### *Xenopus longipes* provides a model to examine the effects of large genome size

Interspecies comparisons of Pipid frogs have provided a powerful approach to characterize scaling relationships and molecular mechanisms of size control^16^. Studies have focused mainly on allotetraploid *X. laevis* (6.35 pg of DNA per nucleus) and diploid *X. tropicalis* (3.6 pg of DNA per nucleus), which possess larger and smaller genome, egg, tadpole, and adult body sizes, respectively^16,18^. However, other related species vary widely in morphometrics and genome content, providing a means to investigate the influence of different size parameters on embryogenesis and potential evolutionary constraints. At one extreme is *Xenopus longipes,* a small frog endemic to one high-altitude lake in Cameroon, Africa (Figure 1A) that has a dodecaploid genome (16 pg of DNA per nucleus), the largest in the *Xenopus* genus^15,17,19^. Its large feet relative to body size distinguishes it from other *Xenopus* species^15,20^ (Figures 1B, S1A and S2A).

**Figure 1.**
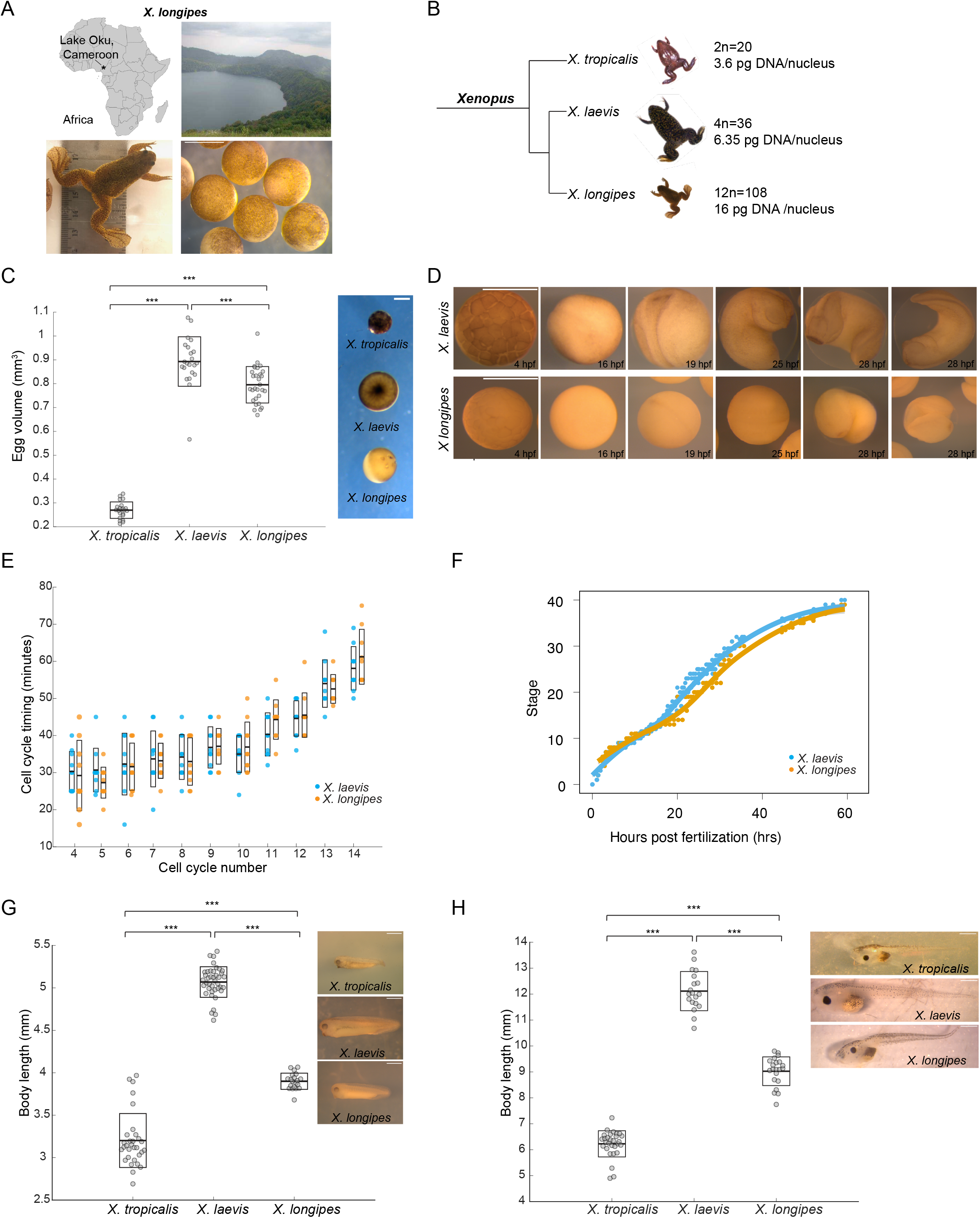
*X. longipes* morphometries and development compared to other *Xenopus* species. (**A**) Schematic of *X. longipes* frog and embryo, and Lake Oku in Cameroon, Africa. Image of Africa from https://en.wikipedia.org/wiki/Africa, Lake Oku picture from Doherty-Bone, 2011. Scale bar 1 mm. (**B**) Phylogeny of *X. longipes* compared to *X. laevis* and *X. tropicalis* with chromosome number and nuclear DNA content. Also see Fig. S1A for detailed frog body length comparison. (**C**) Calculated egg volume in *X. tropicalis, X. laevis,* and *X. longipes*. n≥23 eggs measured in each species. Scale bar, 0.5 mm. (**D**) Images of development in *X. longipes* vs *X. laevis* at 23 ° C. Hpf = hours postfertilization. Scale bar, 1mm. (**E**) Comparison of cell cycle timing in *X. longipes* vs *X. laevis* through the first 14 cleavage divisions. Each dot represents the average timing for 3 individual cells. Thick line inside box = average time, upper and lower box boundaries = +/-std dev. n=3 cells from 10 total embryos analyzed per species from 3 separate clutches. P>0.5 between species in each cell cycle, determined by one-way ANOVA. (**F**) Developmental time course in *X. longipes* vs *X. laevis* at 23° C, 0-62 hpf. n=3 clutches analyzed for *X. laevis,* n=2 for *X. longipes,* 1-3 replicates for each clutch. Also see Video S1, S2, and Fig. S1B for individual timepoints. (**G**) Body length quantification of stage 36 tailbud embryos in *X. tropicalis, X. laevis,* and *X. longipes*. n≥20 tadpoles from 3 clutches measured for each species. Scale bar, 1 mm. (**H**) Body length quantification of stage 48 tadpoles in *X. tropicalis, X. laevis,* and *X. longipes*. n≥20 tadpoles from 3 separate clutches measured for each species. Scale bar, 1 mm. For all box plots, thick line inside box= average length, upper and lower box boundaries= +/- std dev. ***p<0.001, determined by one-way ANOVA.

Despite its large genome, *X. longipes* eggs are slightly smaller than those of *X. laevis* at 1.1 mm in diameter (Figure 1C). We used recently developed husbandry techniques^17^ to generate *X. longipes* embryos via natural matings (Figure 1A), and documented their development compared to *X. laevis* side-by side at the same temperature (Figure 1D). Early development in *X. laevis* has been extensively characterized through metamorphosis, as 12 synchronous and rapid early cleavage divisions generate thousands of individual blastomeres up to the mid-blastula transition (MBT) at Nieuwkoop-Faber stage 8-9, 5-7 hours post fertilization (hpf)^21–23^. *X. longipes* early development proceeded almost indistinguishably through the MBT, but slowed down at neurulation (stage 13, ~16 hpf). Whereas by ~25 hpf, *X. laevis* embryos assumed a tailbud shape with a clear head, tail, and eye protrusions, *X. longipes* embryos remained round, with a closed neural tube but notable lack of anatomical structures (Figure 1D; Video S1). Previous work has documented an inverse correlation between genome size and developmental rate in many organisms^7^,^10^, and studies in frogs indicated longer larval periods in species with larger genomes^8,9,11,24^, in some cases with a direct relationship between amphibian genome size and duration of mitotic and meiotic cell cycles^2,25,26^. However, our analysis revealed only minor changes in developmental timing of early cleavages (Figure 1E), and despite variation in morphology at the late neurula stage 21 (22.5 hpf), cell proliferation was not significantly different between species (Figure S1C). Furthermore, the timing of blastopore closure, which signifies the end of gastrulation^27^, was not delayed in *X. longipes* compared *to X. laevis* (Video S2). During later tailbud stages 35-38 (~50-60 hpf), the delay was much less evident (Figures 1F and 1G) and by the swimming tadpole stage (stage 48, 7 days post fertilization), there were was little discernible difference between the species except for tadpole size (Figure 1H). Thus, while distinct morphological differences were clearly evident at certain developmental timepoints in *X. longipes*, the overall early developmental rate compared to *X. laevis* was similar despite a nearly three-fold difference in genome size.

### Embryo cell size scales with egg size, not genome size, until late in development

In contrast to *X. laevis* and *X. tropicalis*, whose egg and adult body size scales with genome size, the large genome size yet relatively small egg and body size of *X. longipes* provides an extreme counterpoint. Across the tree of life, genome size correlates strongly and linearly with cell and nuclear size^1–6^. We confirmed this conserved relationship in adult frogs ranging nearly 5-fold in ploidy by measuring two cell types in four *Xenopus* species as well as *Hymenochirus boettgeri*, a related Pipid frog. Both erythrocytes (which are nucleated in amphibians) and skin epithelial cells showed a strong, positive, and linear correlation among genome, cell, and nuclear sizes (Figures 2 and S2). Thus, despite their small size relative to other *Xenopus* species, *X. longipes* adults possess large cells corresponding to their high ploidy.

**Figure 2.**
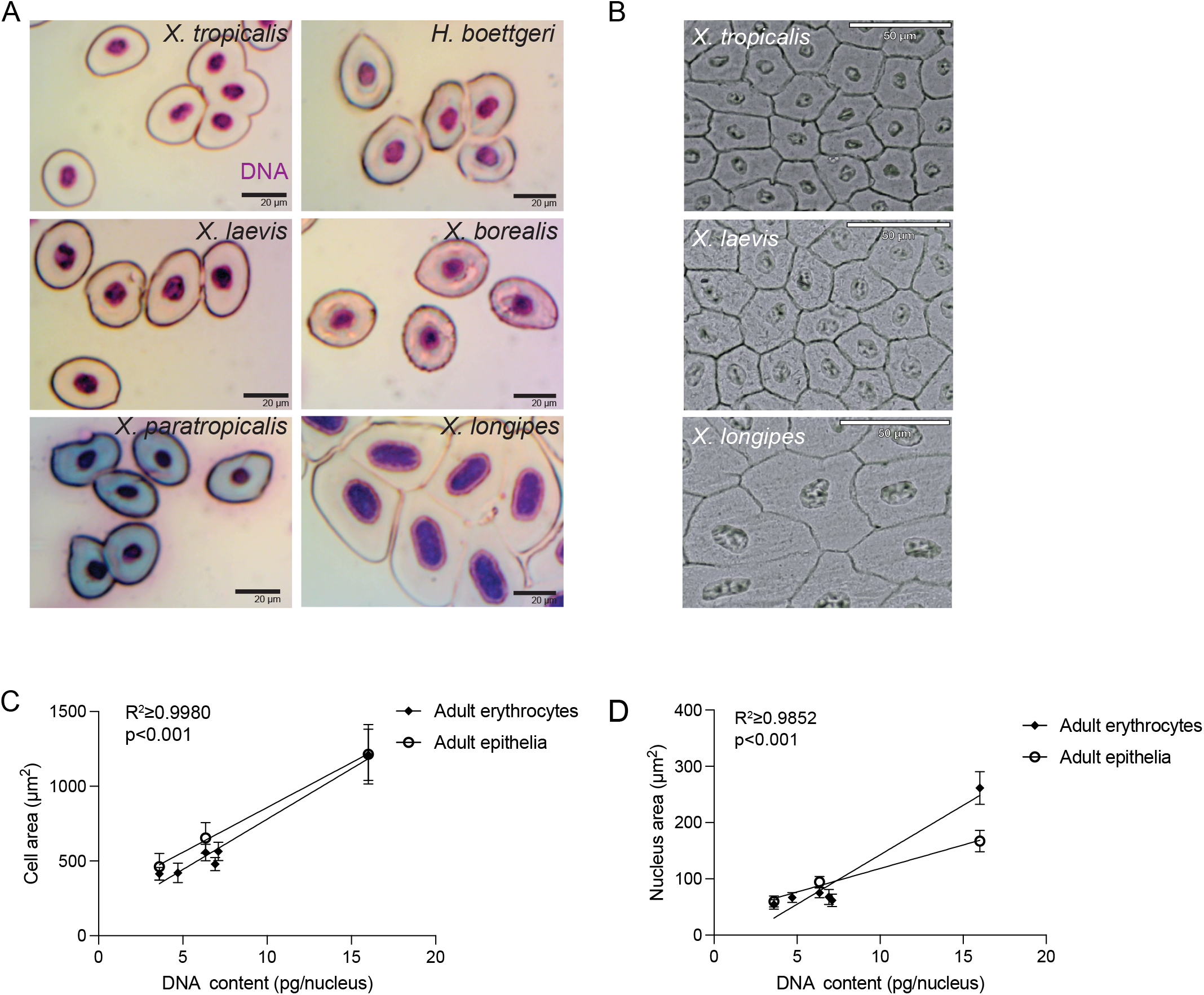
Scaling of cells and nuclei in adult Pipid frog species. (**A**) Images of erythrocytes from 5 adult Pipid frog species of varying ploidies. See Fig. S2A for details about species, including DNA content. Scale bar, 20 μm. (**B**) Images of adult epithelial cells from *X. tropicalis, X. laevis*, and *X. longipes*. Scale bar, 50 μm. (**C**) Average cell cross-sectional area in adult erythrocytes and epithelia plotted as a function of genome size for each frog species. 5 Pipid frog species are plotted for erythrocytes, *X. tropicalis, X. laevis*, and *X. longipes* are plotted for epithelial cells. R^2^ ≥0.9980 for both erythrocytes and epithelia. See Fig. S2B-C for distributions and Fig. S3B for correlation and slope coefficients of each trendline. (**D**) Average nucleus cross-sectional area plotted as a function of genome size for each frog species. 5 Pipid frog species are plotted for erythrocytes, *X. tropicalis*, *X. laevis*, and *X. longipes* are plotted for epithelial cells. R^2^ ≥0.9852 for both erythrocytes and epithelia. See Fig. S2D-E for distributions and Fig. S3E for correlation and slope coefficients of each trendline. For plots in C and D, Error bars= +/- std dev.

Since *X. longipes* frogs possess fewer, larger cells than other *Xenopus* species, we hypothesized that genome to cell size scaling relationships were established during embryogenesis. However, measurements over the course of development (Figures 3A-C) revealed no correlation between cell area and DNA content across species at stage 8, a weak correlation at stages 21 and 36, and a linear relationship only at tadpole stage 48, 7 days post fertilization, when the unfed tadpoles were swimming and possessed well-developed organ and sensory systems^23^ (Figures 3D, S3A and S3B). Instead, cell size varied with egg size throughout development and was strongly, linearly correlated from stage 21 through 36 and persisted through stage 48 (Figures 3E, S3C). In contrast, nuclear size correlated with genome size throughout development (Figures 3A, 3B, S3D, 3F, S3E). Thus, egg volume influences cell size more than genome size between species and genome to cell size scaling does not emerge until quite late in *Xenopus* development, while nuclear size continuously reflects genome size (3G), consistent with a biophysical effect of DNA content on nuclear size^28^.

**Figure 3.**
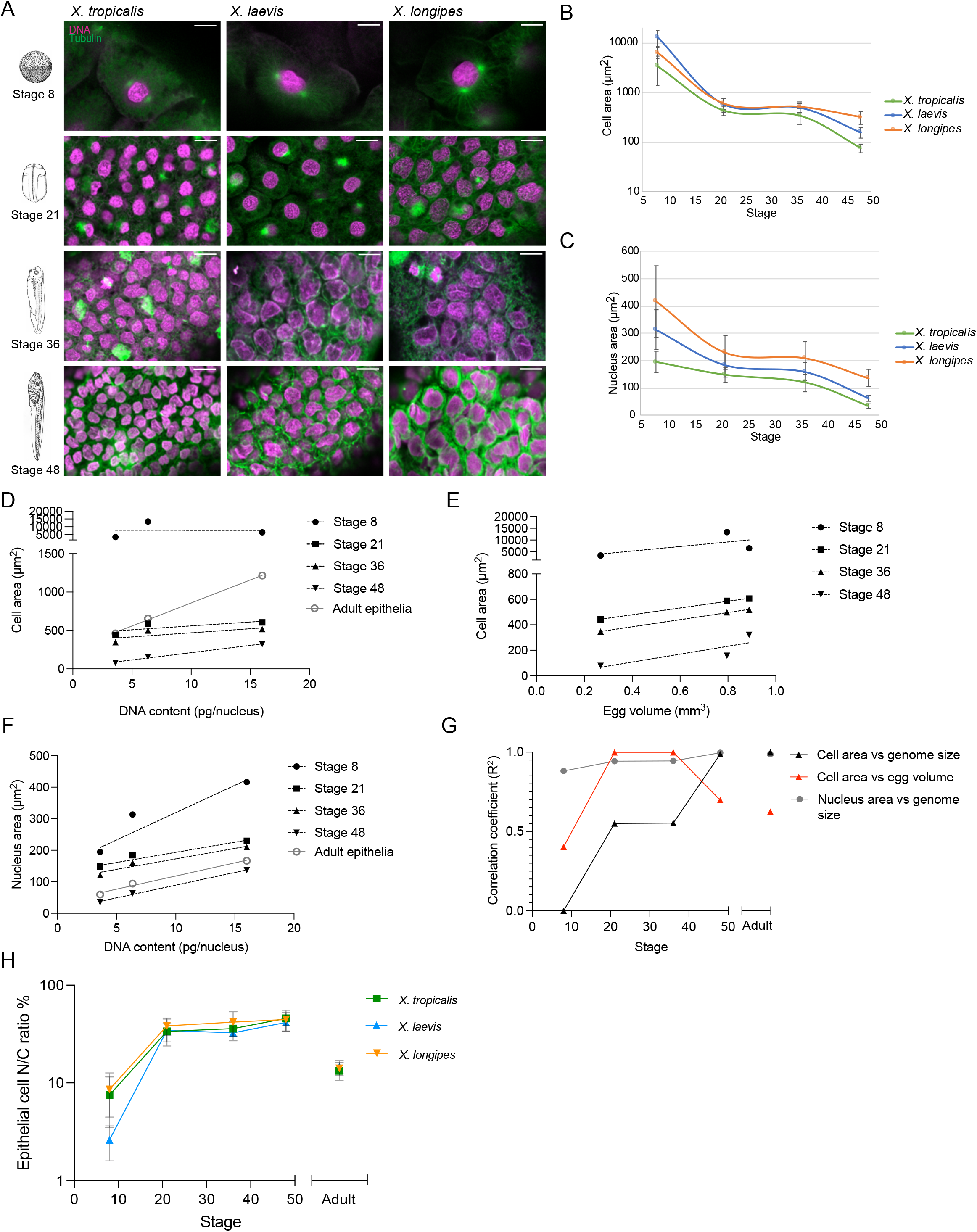
Scaling of cells and nuclei in *Xenopus* embryos. (**A**) Images of cells and nuclei in *X. tropicalis, X. laevis*, and *X. longipes* embryos, stage 8-48. Scale bars, 15 μm. (**B**) Average cross-sectional cell area during embryogenesis, Y-axis plotted in log10. See Fig. S3A for distributions. (**C**) Average cross-sectional nucleus area during embryogenesis. See Fig. S3D for distributions. For plots in B-C, error bars= +/- std. (**D**) Average skin epithelial cell cross-sectional area in embryos and adults plotted as a function of genome size in *X. tropicalis, X. laevis*, and *X. longipes*. See Fig. S3B for correlation and slope coefficients of each trendline. (**E**) Average skin epithelial cell cross-sectional cell area in *X. tropicalis, X. laevis*, and *X. longipes* through embryogenesis, plotted as function of egg volume. See Fig. S3C for correlation and slope coefficients of each trendline. (**F**) Average nucleus cross-sectional area in embryos and adults plotted as a function of genome size in *X. tropicalis*, *X. laevis*, and *X. longipes*. See Fig. S3D for distributions and Fig. S3E for correlation and slope coefficients of each trendline. (**G**) Summary of correlation coefficients of cell or nucleus area vs genome size or egg volume in embryos and adults from Fig. 3D-F, plotted by developmental stage. (**H**) Skin epithelial cell nuclear and cell volumes in embryos and adults of each species were extrapolated from cross-sectional area measurements (Levy et al 2010, Good et al 2013, Jevtic et al 2015), and the ratio of nuclear to cell (N/C) volume expressed as a percentage. Average values are plotted, Y-axis plotted in log_10_. Error bars= +/- std. Also see Fig. S3F for distribution of N/C ratios.

### Embryogenesis is characterized by distinct size scaling regimes

Another way to evaluate changing size relationships across embryos of different *Xenopus* species is to characterize the nuclear-cytoplasmic (N/C) ratio, which is precisely maintained in eukaryotic organisms and differentiated cell types^13^. During development, a threshold N/C value is thought to trigger the MBT when rapid cleavages abruptly cease, cycles of slow, asynchronous divisions begin, the zygotic genome is activated, and gastrulation and cell differentiation initiates^22,29^. MBT timing is influenced by both ploidy and cell size within a species^22,30,31^. However, measurements comparing embryos from the three different frogs upon MBT onset at stage 8 revealed that *X. laevis* cells possess a significantly lower N/C ratio than either *X. tropicalis* or *X. longipes* (Figure 3H, S3F). At this point in development there was no correlation between genome size and cell size (Figure 3D). These findings suggest that MBT timing is triggered at different N/C ratios in different frog species. Current models propose that titration of nuclear or chromatin factors trigger the MBT^32,33^. Future experiments will elucidate whether maternal supplies are tuned according to egg and genome sizes so that the same number of cleavage divisions leads to similar MBT timing despite very different size metrics.

Beyond the MBT, between stages 21 and 36 of development that includes neurulation, the N/C ratio was similar in epithelial cells of all three *Xenopus* species (Figure 3H). During this period, rapid cell divisions have ceased and cell and nuclear sizes remain relatively constant (Figures 3B and 3C). Between stages 36 and 48, cell sizes again decreased, but now correlated with genome size, with *X. tropicalis* cells becoming smaller than *X. laevis* cells, which in turn were smaller than *X. longipes* cells, reflecting the emergence of genome to cell size scaling (Figures 3B and 3C). Nuclear sizes also decreased during this phase so that N/C ratios remained constant. However, embryo epithelial cells at stage 48 were smaller than adult skin cells across all three species, while nuclear sizes between adults and tadpoles were similar, leading to a significantly lower N/C ratio in adult epithelia, which was similar to that of adult erythrocytes (Figures 3B, 3C, 3H and S3F). Therefore, whereas cell size and N/C ratio do not match adult dimensions in swimming tadpoles, nuclear size does.

Altogether, these results indicate that distinct scaling regimes exist during development that may reflect the physiology of the embryo. In a first regime, embryos initially undergo rapid cleavage divisions that increase cell number and decrease cell size exponentially. Genome size during this regime appears to be irrelevant as cell cycle timing of *X. longipes* was indistinguishable from *X. laevis* (Figure 1E). A second regime appears once a stable N/C ratio is established by neurulation at stage 21. During this period, complex cell movements are coordinated with differentiation and cell divisions occur at a much lower frequency^34^. In a third regime, N/C ratios remain constant but genome to cell size scaling is established by stage 48. Finally, larger, adult cell sizes and correspondingly lower N/C ratios emerge, which coincides with termination of the maternal program and the onset of tadpole feeding.

### Embryo nuclei and spindles scale with genome and cell size, respectively

Scaling of subcellular structures including the nucleus and the spindle to cell size has been characterized across *Xenopus* species and during early development, and molecular mechanisms identified that coordinately scale both structures according to cell surface-to-volume ratio^35^. Although nuclear and spindle size has been documented extensively in *Xenopus* and other organisms in early cleaving embryos prior to the MBT^35–39^, much less is known about subcellular scaling later in embryonic development following the onset of zygotic transcription and morphogenesis. Our analysis revealed that although nuclear size correlated with cell size as widely reported across species and cell types^13^, it was more highly correlated with genome size throughout development, starting as early as stage 8 (Figures 4A, 4B, S4A and S4B). In contrast, we found that spindle length was only moderately correlated with genome size between stage 8 and 36, consistent with previous findings^38,39^ (Figures 4C, S4C, 4D and S4B). Instead, spindle sized was strongly and linearly correlated with egg volume and cell size (Figures 4E–4G and S4D). Thus, nuclei and spindles possess distinct scaling properties through development, with nuclear size influenced more by genome size and spindle size by cell size.

**Figure 4.**
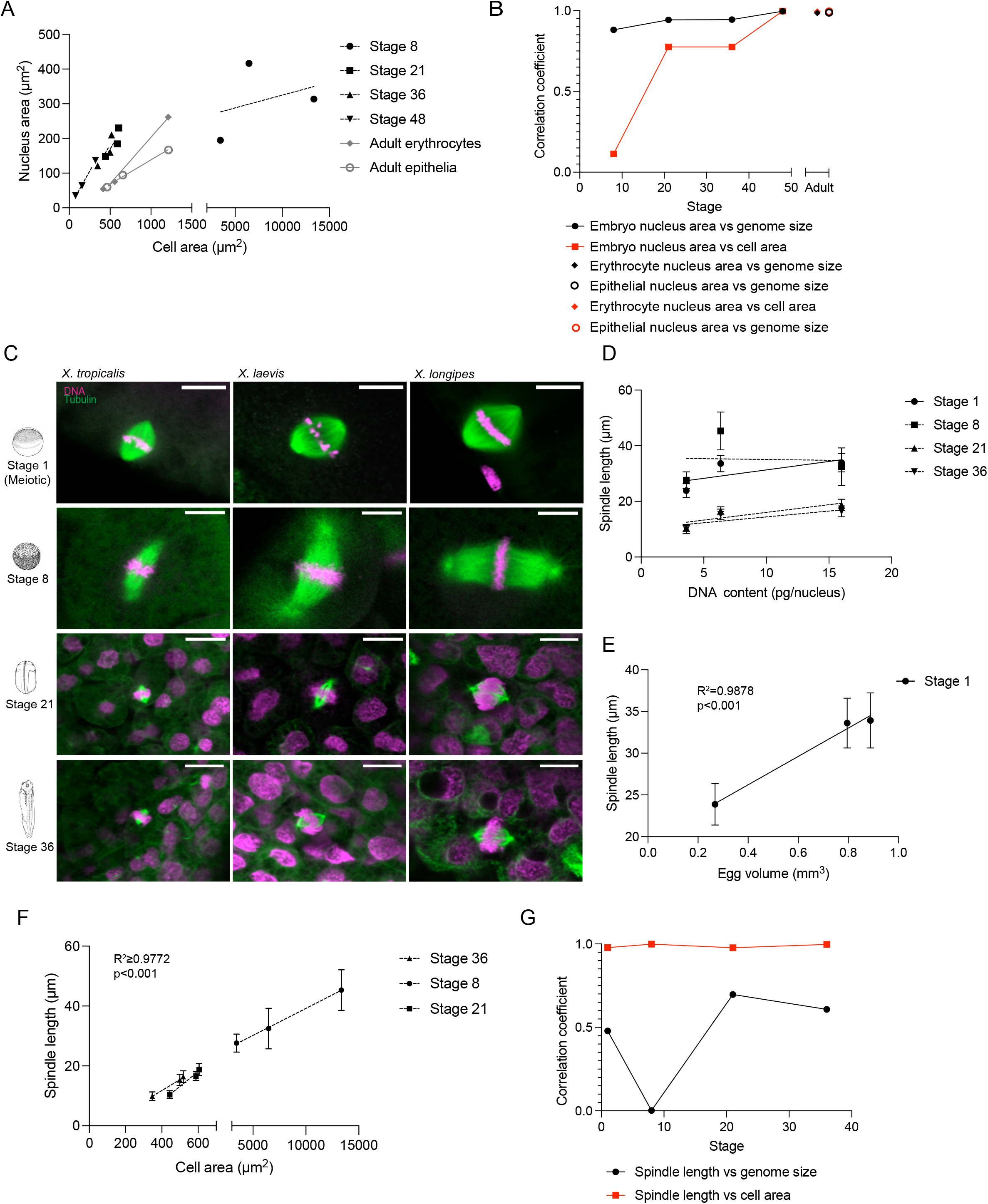
Subcellular scaling of nuclei and spindles through development. (**A**) Average cross-sectional nucleus area in *X. tropicalis, X. laevis*, and *X. longipes* embryos and adults, plotted as function of cell area. (**B**) Summary of correlation coefficients of nucleus area vs genome size and cell area through development from Figure 3F and 4A. (**C**) Images of meiotic and mitotic spindles in *X. tropicalis, X. laevis*, and *X. longipes* embryos, stage 1 (unfertilized meiotic) through stage 36. Scale bars= 20 μm. (**D**) Spindle length plotted as a function of genome size in *X. tropicalis, X. laevis*, and *X. longipes* embryos. See Fig. S4B for correlation and slope coefficients of each trendline, and Fig. S4C for distributions. (**E**) Egg spindle length (stage 1, meiotic) plotted as a function of egg volume. See Figure S4C for distribution. (**F**) Average spindle length plotted as a function of cell area through development in *X. tropicalis, X. laevis*, and *X. longipes* embryos. See Figure S4C for distribution, and S4D for correlation and slope coefficients of each trendline. (**G**) Summary of correlation coefficients of spindle length vs genome size or cell area through development from Figure 4D-F. For all plots, error bars = +/- std dev.

In conclusion, the gradual emergence of genome to cell size scaling we observed in different *Xenopus* species during development reveals that the cell division cycle does not acutely adapt to genome size, and that the transition to adult size scaling relationships (at least in skin epithelia) occurs after tadpoles begin feeding. Instead, egg size strongly influences cell size scaling during embryogenesis, highlighting the key role of maternally contributed components to the developmental program^40^. Interestingly, across *Xenopus* species MBT occurs with similar timing despite very different size metrics. Thus, over tens of millions of years of evolution, the basic developmental program in *Xenopus* frogs has remained robust to a six-fold change in genome size. Whether the observed developmental delay at neurulation and differences in *X*. *longipes* embryo morphogenesis result from its large genome is an open question, since cell sizes during this period are very similar to *X. laevis*, although nuclear sizes and N/C ratios are slightly larger in surface epithelia. Further characterization of *X. longipes* embryogenesis and gene expression patterns will be informative in understanding the basis of this variation.

Our inter-species comparison indicates that different scaling regimes operate during development. While the molecular mechanisms that underlie cell size scaling remain mysterious, future work will reveal how different size-dependent modalities emerge based on maternal resource allocation and temporal activation of growth signaling pathways, which in turn promise to shed light on when and how single-cell metabolic and biosynthesis properties, which correlate with genome size^41–43^, affect tissue and whole organism physiology. In addition to serving as an extreme case for investigating genome size effects on development and physiology, *X. longipes* also provides a powerful model to study adaptations at the subcellular level. Whether and how 108 chromosomes impact nuclear and spindle organization, as well as the process of cell division, can be investigated in embryos and also using powerful in vitro systems unique to *Xenopus*^16^.

## ACKNOWLEDGEMENTS

We thank Nicole Chaney and California Academy of Sciences for *X. longipes* frogs, as well as valuable husbandry advice, Ben Evans for helpful advice and *X. paratropicalis* blood smears, Jim Evans and Mike Fitzsimmons for dedicated and careful tadpole rearing, Scott Aposhian for initial measurements of erythrocytes, Helena Cantwell for critical reading of the manuscript, and members of the Heald lab, past and present, for helpful discussions and support with experiments. R.H. was supported by NIH MIRA grant R35 GM118183 and the Flora Lamson Hewlett Chair.

## AUTHOR CONTRIBUTIONS

Conceptualization, Funding acquisition: RH, KM, CC

Methodology, Investigation, Visualization: KM, CC

Supervision: RH

Manuscript preparation: KM, RH, CC

## SUPPLEMENTARY MATERIALS

Materials and Methods

Figures S1 to S4

Videos S1 and S2

## MATERIALS AND METHODS

### Erythrocyte preparation and measurements

A small drop of blood was collected from the frog foot of each species with a sterile needle, and the drop was smeared on a slide. The smear was then fixed with methanol and stained with Giemsa stain (Sigma GS). Cells were imaged in brightfield using micromanager software ^44^ with an upright Olympus BX51 microscope equipped with a Olympus UPlan 40x air objective and ORCA-II camera (Hamamatsu Photonics, Hamamatsu city, Japan). Cross-sectional areas of cells and nuclei were measured in Fiji using the freehand tool. *X. paratropicalis* blood smears were a kind gift from Ben Evans (McMaster University).

### Epithelial cell preparation and measurements

Shed frog skin was collected from frog housing tanks and mounted carefully on a microscope slide by rolling the skin flat so that a monolayer of cells could be imaged. Cells were imaged immediately after collection, unstained and without a coverslip, in brightfield using Olympus cellSens Dimension 2 software on an upright Olympus BX51 microscope equipped with an ORCA-II camera (Hamamatsu Photonics) and an Olympus UPlan 20x air objective. Cross sectional areas of cells and nuclei were measured in Fiji using the freehand tool.

### Natural mating of *X. longipes*

*X. longipes* were a kind gift from California Academy of Sciences. Male and female *X. longipes* were injected with a priming dose of 75 IU HCG (Sigma) 48 hours before the desired mating day, and kept separately to avoid premature amplexus. On the day of ovulation, males and females were injected with a boosting dose of 200 IU HCG. Amplexus began soon after injection with egg laying 6-8 hours later. Embryos were collected in batches for fixation and live imaging.

### In vitro fertilization of *X. laevis*

*X. laevis* females were primed with 100 IU of pregnant mare serum gonadotropin (PMSG, National Hormone and Peptide Program, Torrance, CA) at least 48 h before use and boosted with 500 IU of HCG (Human Chorionic Gonadotropin CG10, Sigma) 14-16 hours before experiments. To obtain testes, males were euthanized by anesthesia through immersion in double-distilled (dd)H2O containing 0.15% MS222 (tricaine) neutralized with 5 mM sodium bicarbonate before dissection. Testes were collected in 1X Modified Ringer (MR) (100 mM NaCl, 1.8 mM KCl, 1 mM MgCl2, 5 mM HEPES-NaOH pH 7.6 in ddH2O) and stored at 4°C until fertilization. To prepare the sperm solution, 1/3 testis was added to 1 mL of ddH2O in a 1.5 mL microcentrifuge tube, and homogenized using scissors and a pestle. *X. laevis* females were squeezed gently to deposit eggs onto petri dishes coated with 1.5% agarose in 1/10X MMR. Any liquid in the petri dishes was removed and the eggs were fertilized with 1 mL of sperm solution per dish. Fertilized embryos were swirled in the solution to form a monolayer on the bottom of the petri dish and incubated for 10 min with the dish slanted to ensure submersion of eggs. Dishes were then flooded with 1/10X MMR, swirled and incubated for 10 min. To remove egg jelly coats, the 1/10X MMR was completely exchanged for freshly prepared Dejellying Solution (2% L-cysteine in ddH2O-NaOH, pH 7.8). After dejellying, eggs were washed extensively (>4X) with 1/10X MMR before incubation at 23°C. At Nieuwkoop and Faber stage 2-3, fertilized embryos were sorted and placed in fresh 1/10X MMR in new petri dishes coated with 1.5% agarose in 1/10X MMR.

### Maintenance of *Xenopus* embryos

*X. laevis, tropicalis*, and *longipes* embryos were raised side by side in 1.5% agarose in 1/10X MMR -coated petri dishes covered in 1/10X MMR in a 23 °C incubator. The MMR was changed and dead/lysed embryos removed frequently to prevent contamination.

### Imaging and measurement of egg diameters, developing embryos, and tadpoles

For still images, eggs and embryos were placed in an agarose-coated imaging chamber filled with a limited amount of 1/10X MMR to prevent depth-biased measurements and imaged at 12x-31x magnification using a Wild Heerbrugg M7A StereoZoom microscope coupled to a Leica MC170HD camera and Leica LAS X software. Diameter of eggs was measured using the line tool in Fiji.

### Live imaging

Movies of *Xenopus* embryo development were made by placing embryos in 1/10X MMR in imaging dishes prepared using an in-house PDMS mold designed to create a pattern of 1.5 mm large wells in agarose that allowed us to image embryos of each species simultaneously. Time-lapse movies were acquired either at a frequency of one frame every 10 s for 20 h, or one frame every 4 minutes for x hours. Movies are compressed to 15 frames per second.

### Cell Cycle Duration measurement

Cell cycles were measured as described in^33^. Movies were started at ~2 h post-fertilization (after the first or second cleavage). Embryos were allowed to develop in 1/10X MMR. Divisions were counted to determine the frame number of the eighth cleavage. Then, ~15 individual cells were selected from the visible portion of each embryo after the eighth cleavage. The period between cleavages was determined by manually tracking individual cells and noting the frame number at which the cleavage furrow visibly transected the entire cell. When daughter cells did not divide concurrently, the division time of the earliest dividing daughter was always used, and that cell was followed for the remainder of the movie. When the cleavage could not be observed, as in cases where the cleavage plane did not intersect the embryo surface, the cell was omitted from analysis.

P-values were calculated using a two-tailed t-test with unequal variance between the indicated distributions. At least 3 cells were counted for each embryo at each cell cycle using 10 total embryos from 3 separate clutches.

### Embryo whole mount immunofluorescence

To label nuclei, cell borders, and mitotic/meiotic spindles, embryos at the desired developmental stage were fixed for one hour using MAD fixative (2 parts methanol [Thermo Fisher Scientific, Waltham, MA], 2 parts acetone [Thermo Fisher Scientific,

Waltham, MA]), 1 part DMSO [Sigma]). After fixation, embryos were dehydrated in methanol and stored at −20°C. Embryos were then processed as previously described ^45^ with modifications. Following gradual rehydration in 0.5X SSC (1X SSC: 150 mM NaCl, 15 mM Na citrate, pH 7.0), embryos were bleached with 1-2% H2O2 (Thermo Fisher Scientific, Waltham, MA) in 0.5X SSC containing 5% formamide (Sigma) for 2-3 h under light, then washed in PBT (137 mM NaCl, 2.7 mM KCl, 10 mM Na2HPO4, 0.1% Triton X-100 [Thermo Fisher Scientific, Waltham, MA]) and 2 mg/mL bovine serum albumin (BSA). Embryos were blocked in PBT supplemented with 10% goat serum (Gibco – Thermo Fisher Scientific, Waltham, MA) and 5% DMSO for 1-3 h and incubated overnight at 4°C in PBT supplemented with 10% goat serum and primary antibodies. The following antibodies were used to label tubulin, DNA, and Phospho-histone H3: 1:250 mouse anti-beta tubulin (E7; Developmental Studies Hybridoma Bank, Iowa City, IA) 1:250 rabbit anti-histone H3 (ab1791; Abcam, Cambridge, MA), 1:1000 Anti-phospho-Histone-H3(ser10) (06-570; EMD Millipore). Embryos were then washed 4× 2 h in PBT and incubated overnight in PBT supplemented with 1:500 goat anti-mouse or goat anti-rabbit secondary antibodies coupled either to Alexa Fluor 488 or 568 (Invitrogen – Thermo Fisher Scientific, Waltham, MA). Embryos were then washed 4× 2 h in PBT and gradually dehydrated in methanol. Embryos were cleared in Murray’s clearing medium (2 parts of Benzyl Benzoate, 1 part of Benzyl Alcohol).

### Confocal imaging and measurement of embryos, cells and nuclei after whole mount immunofluorescence

Embryos were placed in a reusable chamber (Thermo Fisher) in fresh Murray’s clearing medium for confocal microscopy. Confocal microscopy was performed on a Zeiss LSM 800 confocal running the Zeiss Zen Software. Embryos were imaged using a Plan-Achromat 20x/0.8 air objective and laser power of 0.5-2%, on multiple 1024×1024 pixel plans spaced 0.68-1.2 μm apart in Z. In stage 8 embryos, before differentiation, we measured an equal distribution of both animal and vegetal cells closest to the surface of the embryo, where staining was most penetrant. In stage 21, 36, and 48 embryos, we measured both ciliated and unciliated cells of the surface epithelium. Cross-sectional areas of cells and nuclei were measured in Fiji at the central plane of the cell or nucleus using the freehand tool.

### Egg volume and N/C ratio calculations

Egg volume was calculated from 2D stereo images of eggs from each species as described above in “Imaging and measurement of egg diameters, developing embryos, and tadpoles”. The diameter of each egg was measured using the line tool in Fiji and the volume calculated using the formula for volume of a sphere (V = 4/3 πr^3^). N/C ratios were extrapolated from 2D cross-sectional area measurements using a method described and validated for embryos ≥stage 8^36,37,39^. The cross-sectional area of the nucleus at its widest part was divided by the cross-sectional area of the cell at the same location in Z, then multiplied by 100 to express N/C volume ratio expressed as a percent.

### Data analysis and statistics

Box plots were generated in Matlab (Figures 1E, 1G, 1H) or using GraphPad Prism software (all others) which plot the mean and standard deviation for each condition. P-values between averages were generated using a one way ANOVA. Simple linear regression plots (ie. In 2C,2D, 2H) were generated using GraphPad Prism software, which calculated the Pearson’s Correlation Coefficient (R), trendline for regression (R^2^), and Slope Coefficient for each trendline. P-values between slopes were calculated using Prism’s pairwise Analysis of Covariance (ANCOVA) test.

---

**Figure S1.**
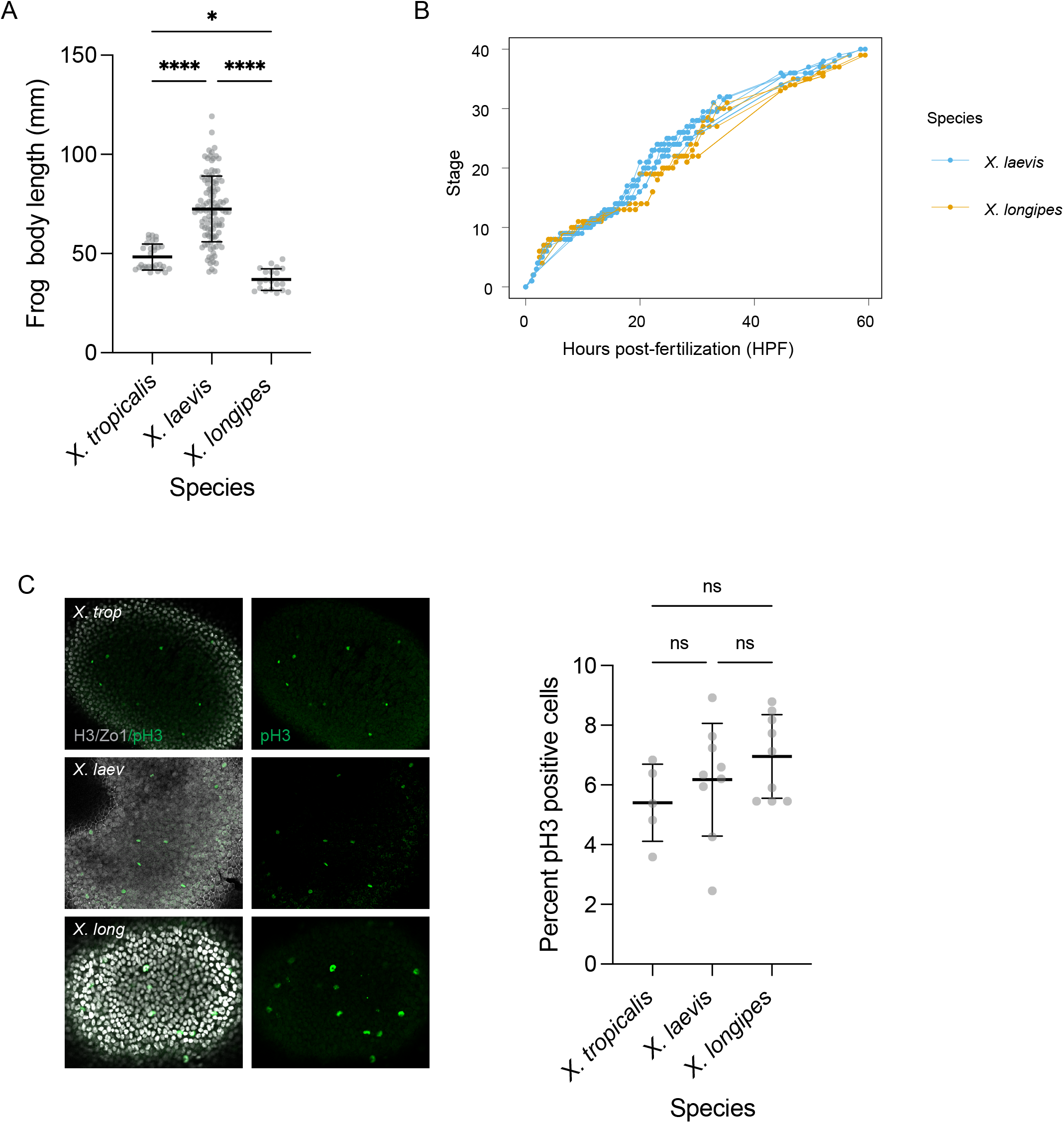
*X. longipes* body length and further development characterization. Related to Figure 1. (**A**) Snout to vent length plotted for male and female adult *X. tropicalis, X. laevis*, and *X. longipes* frogs. n=21 *X. longipes* measured. Lengths of *X. tropicalis* (n=38) and *X. laevis* (n=110) are from Evans et al, 2015. *p<0.05, ***p<0.001, determined by one-way ANOVA. (**B**) Developmental time course in *X. longipes* vs *X. laevis*, without smoothened line, showing individual timepoints. (**C**) Immunofluorescence staining of stage 21 *X. tropicalis, X. laevis*, and *X. longipes* embryos to assess the fraction of mitotic cells in each species. Cell borders (stained for Zo1) and nuclei (stained for histone H3) are shown in gray. Dividing cells are stained with M-phase marker phospho-H3 (pH3). Quantification shows percent of pH3-positive cells in each species. Thick line in the middle of each set of points = mean, error bars= +/- std dev. ns= not significant, determined by one-way ANOVA.

**Figure S2.**
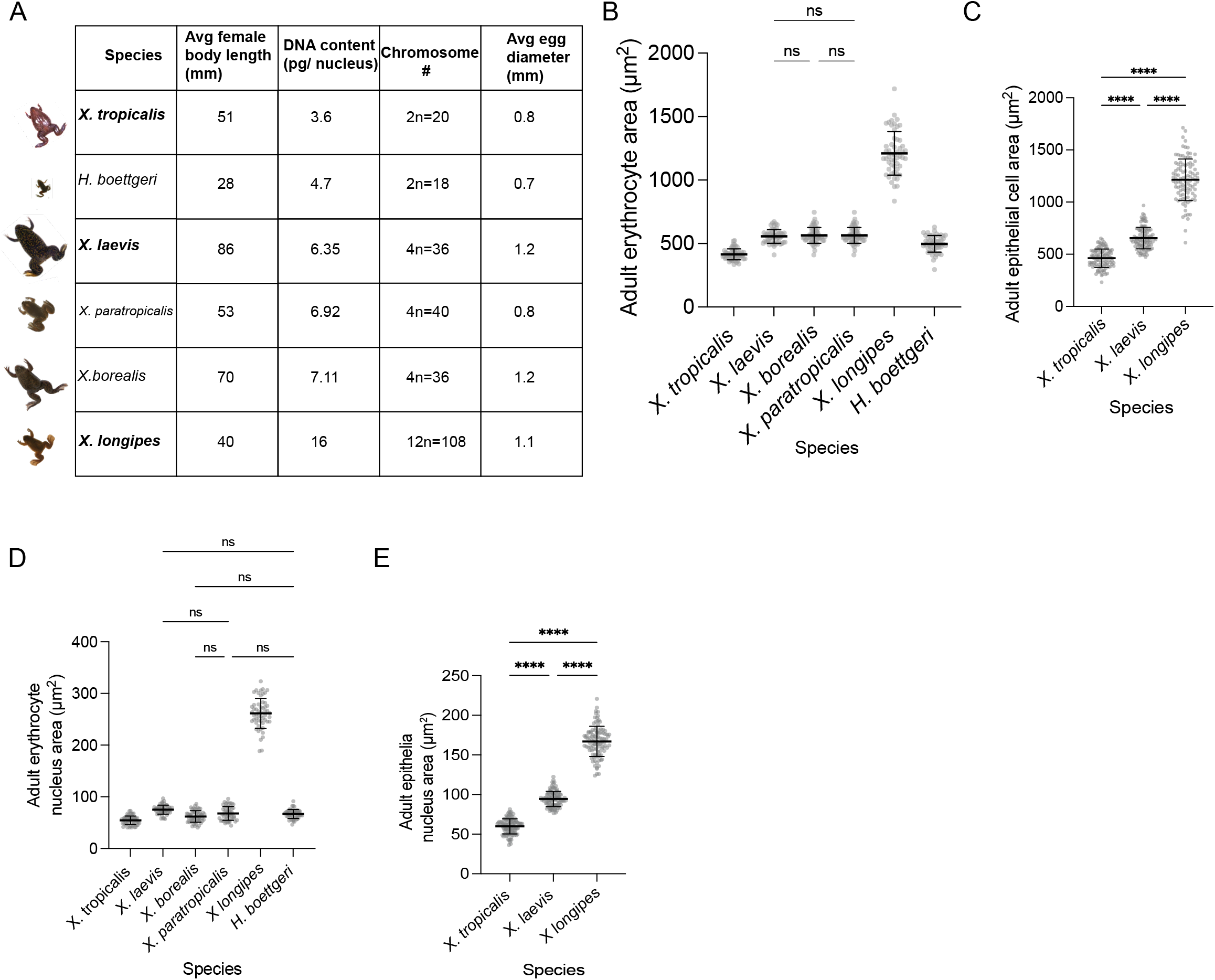
Measurements of cells and nuclei in adult Pipid frog species. Related to Figure 2. (**A**) Characteristics of frog species used for erythrocyte measurements. (**B**) Cross-sectional cell area of erythrocytes from adult frog species described in S1A. All P-values between species are <0.001 (not shown), except where indicated as not significant (ns) determined by one-way ANOVA. (**C**) Cross-sectional cell area of epithelial cells from adult *X. tropicalis, X. longipes*, and *X. laevis*. (**D**) Cross-sectional nucleus area of erythrocytes from adult frog species described in S1A. All p-values between species are <0.001 (not shown), except where indicated as not significant (ns) determined by one-way ANOVA. (**E**) Cross-sectional nucleus area of epithelial cells from adult *X. tropicalis, X. longipes*,and *X. laevis*. For plots in B-E, thick line in the middle of each set of points = mean, error bars = +/- std dev. ***p<0.001. For erythrocytes, n≥50 cells and nuclei were measured in each species. For epithelia, n≥100 cells and nuclei were measured in each species.

**Figure S3.**
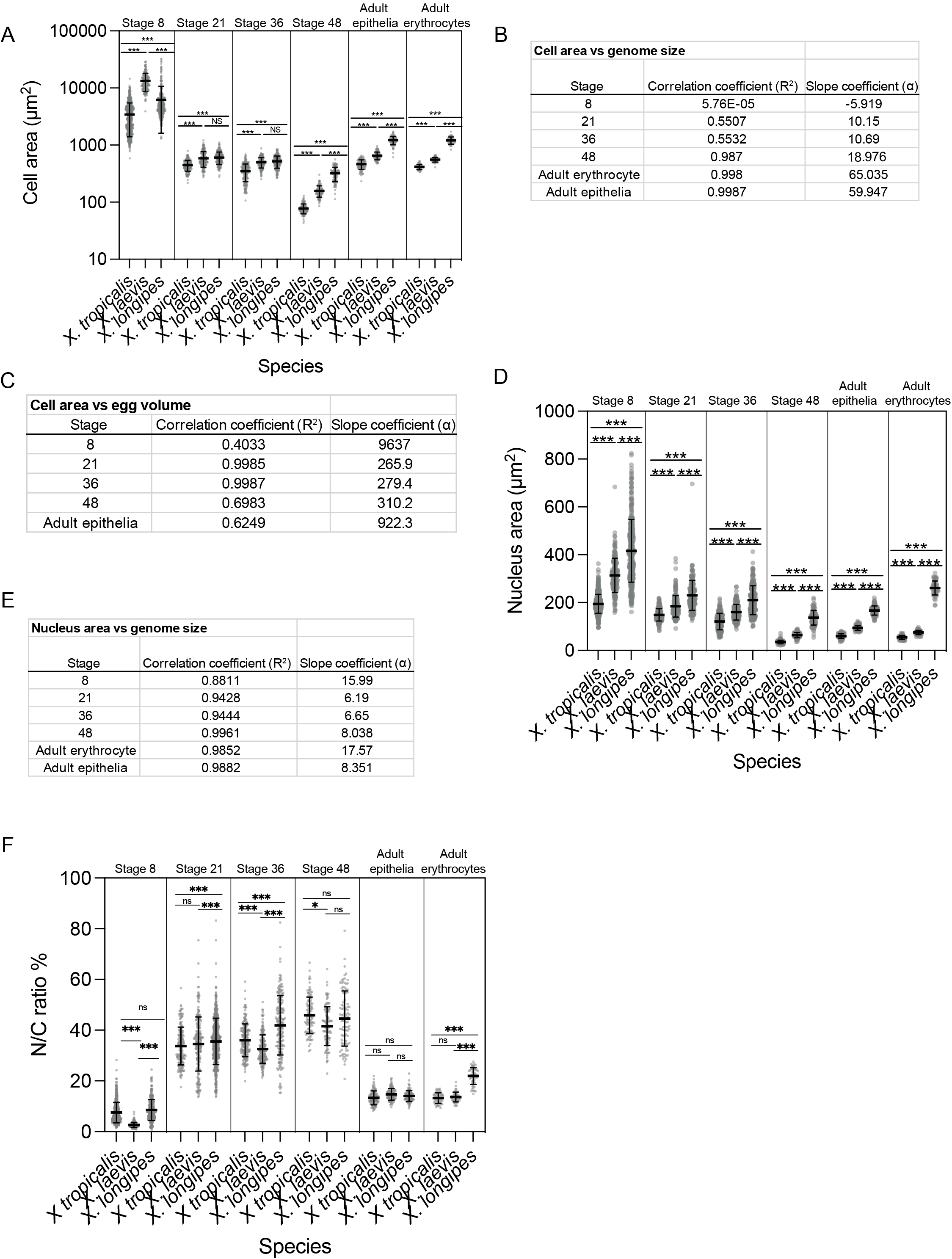
Measurement of cells, nuclei and N/C ratios in *Xenopus* embryos. Related to Figures 2 and 3. (**A**) Cross-sectional areas of cells from *X. laevis, X. tropicalis*, and *X. longipes* embryos and adults. (**B**) Correlation and slope coefficients of cell area vs genome size from linear trendlines in Fig. 3D. Correlation coefficients for linear trendlines (R^2^) were calculated from the Pearson’s Correlation Coefficient (R). To determine whether slope coefficients were significantly different, we calculated P values using a pairwise Analysis of Covariance (ANCOVA). Slope coefficients were statistically similar between stage 21 and 36 (p=0.9713) and different between stage 48 and adult epithelia (p=0.0056). (**C**) Correlation and slope coefficients of cell area vs egg volume from linear trendlines in Fig. 3E. Correlation coefficients for linear trendlines (R^2^) were calculated from the Pearson’s Correlation Coefficient (R). (**D**) Cross-sectional area of nuclei from *X. laevis, X. tropicalis*, and *X. longipes* embryos and adults. (**E**) Correlation and slope coefficients of nucleus area vs genome size from linear trendlines in Fig. 3C. Slope coefficients were statistically similar between stage 21 and 36 (p=0.6134) and between stage 48 and adult epithelia (p=0.2344). (**F**) N/C volume ratios in *X. tropicalis, X. laevis*, and *X. longipes*, embryos and adults, calculated from cross-sectional area measurements in 3B-C. For plots in A, D, and F, at stage 8, cells near the surface of the embryo from both animal and vegetal polls were measured. From stage 21-48, skin epithelial cells were measured. n≥100 cells or nuclei from ≥3 embryos or adults from ≥3 separate clutches were measured per stage for each species. Thick line in the middle of each set of points=mean. For all plots, error bars= +/- std dev. ***p<0.001 and ns= not significant, determined by one-way ANOVA.

**Figure S4.**
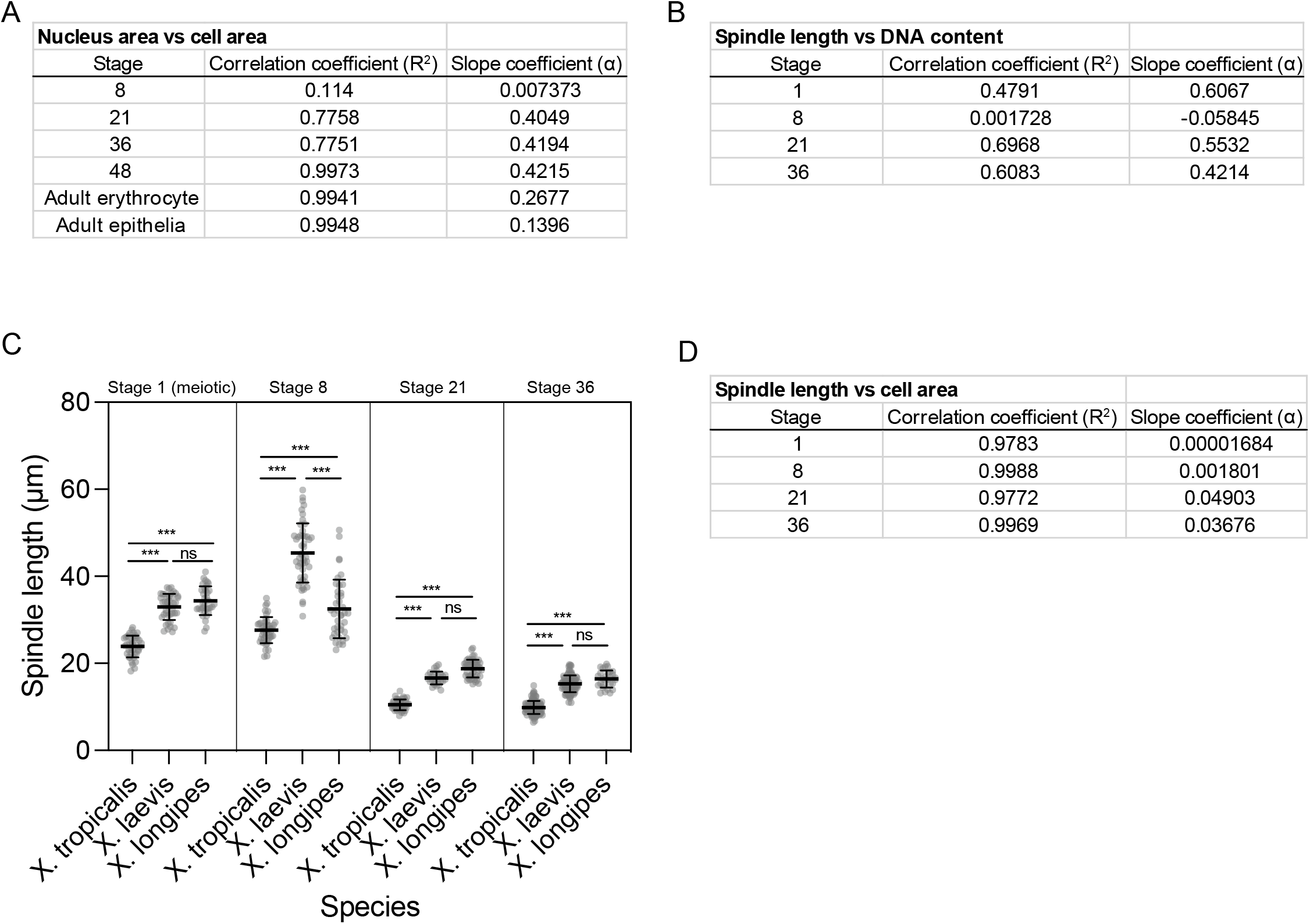
Spindle length in *X. tropicalis, X. laevis*, and *X. longipes* embryos. Related to Figure 3. (**A**) Correlation and slope coefficients of nucleus area vs cell area from linear trendlines in Fig. 4A. Correlation coefficients for linear trendlines (R^2^) were calculated from the Pearson’s Correlation Coefficient (R). (**B**) Correlation and slope coefficients of spindle length vs genome size from linear trendlines in Fig. 4D. (**C**) Spindle length (measured as pole-to-pole distance) through development including Stage 1 (unfertilized meiotic). Thick line in the middle of each set of points=mean. Error bars= +/- std dev. ***p<0.001, ns= not significant, determined by one-way ANOVA. (**D**) Correlation and slope coefficients of spindle length vs cell area from linear trendlines in Fig. 4F. n≥34 spindles from >3 eggs/embryos from ≥3 clutches measured for each species per stage.

**Video S1. Related to Figure 1.** Comparison of early development in *X. longipes* (right) *X. laevis* (left) from animal pole, early cleavage through tailbud stage.

**Video S2. Related to Figure 1.** Comparison of early development in *X. longipes* (left) and *X. laevis* (right) from vegetal pole, early cleavage through neurula stage.

